# Cryo-EM Reveals Ligand Induced Allostery Underlying InsP_3_R Channel Gating

**DOI:** 10.1101/374041

**Authors:** Guizhen Fan, Mariah R. Baker, Zhao Wang, Alexander B. Seryshev, Steven J. Ludtke, Matthew L. Baker, Irina I. Serysheva

**Affiliations:** Department of Biochemistry and Molecular Biology, Structural Biology Imaging Center, McGovern Medical School at The University of Texas Health Science Center at Houston, 6431 Fannin Street, Houston, Texas 77030, USA; Verna and Marrs McLean Department of Biochemistry and Molecular Biology, Baylor College of Medicine, One Baylor Plaza, Houston, Texas 77030, USA

## Abstract

Inositol-1,4,5-trisphosphate receptors (InsP_3_Rs) are cation channels that mobilize Ca^2+^ from intracellular stores in response to a wide range of cellular stimuli. The paradigm of InsP_3_R activation is the coupled interplay between binding of InsP_3_ and Ca^2+^ that switches the ion conduction pathway between closed and open states to enable the passage of Ca^2+^ through the channel. However, the molecular mechanism of how the receptor senses and decodes ligand-binding signals into gating motion remains unknown. Here we present the electron cryo-microscopy structure of InsP_3_R1 from rat cerebellum determined to 4.1 Å resolution in the presence of activating concentrations of Ca^2+^ and adenophostin A (AdA), a structural mimetic of InsP_3_ and the most potent known agonist of the channel. Comparison with the 3.9 Å-resolution structure of InsP_3_R1 in the Apo-state, also reported herein, reveals the binding arrangement of AdA in the tetrameric channel assembly and striking ligand-induced conformational rearrangements within cytoplasmic domains coupled to the dilation of a hydrophobic constriction at the gate. Together, our results provide critical insights into the mechanistic principles by which ligand-binding allosterically gates InsP_3_R channel.

## Introduction

Inositol 1,4,5-trisphosphate receptors (InsP_3_Rs) constitute a functionally important class of intracellular Ca^2+^ channels that are capable of converting a wide variety of external and internal signals received by the cell (e.g. hormones, neurotransmitters, growth factors, light, odorants) to intracellular calcium signals, which trigger markedly different cellular actions ranging from gene transcription to secretion, from proliferation to cell death^1^. The cellular signals are transmitted to the receptor by the secondary messenger molecule inositol 1,4,5-trisphosphate (InsP_3_), the primary agonist of InsP_3_Rs, generated within an essential intracellular signaling pathway initiated by phospholipase C. There is a general consensus that activation of channel gating is associated with conformational rearrangements at the inner pore-lining helix bundle that are triggered by InsP_3_ binding within the first 600 residues of the InsP_3_R protein^2,3^. This functional coupling has been experimentally demonstrated through electrophysiological, ligand-binding and mutagenesis studies^4,5^, however the precise molecular mechanism by which InsP_3_ exerts its effect on InsP_3_R function is largely unknown. Our previous study described the 4.7 Å resolution electron cryomicroscopy (cryo-EM) structure of the full-length tetrameric InsP_3_R1 channel in an Apo-state, which revealed a network of intra- and inter-domain interfaces that might be responsible for the conformational coupling between ligand-binding and gating activation^2^. To further investigate how the structure of the InsP_3_R channel allows for ligand initiated gating we have now determined the 3D structure of InsP_3_R1 bound to adenophostin A (AdA), a highly potent agonist of InsP_3_Rs^6,7^, to 4.1 Å resolution using single-particle cryo-EM analysis. In this study, we have also extended our structural analysis of InsP_3_R1 in a ligand-free (Apo) state to 3.9 Å resolution. Together, these structures reveal how InsP_3_R1 channel performs its mechanical work through ligand-driven allostery that alleviates the molecular barrier within the ion permeation pathway and allows for Ca^2+^ translocation across the membrane.

## Results

### Structure of AdA-InsP_3_R1

To understand how ligand-binding triggers a drastic change in the permeability of InsP_3_R channel to specific ions, we determined the structure of InsP_3_R1 in the presence of activating concentrations of AdA (100 nM) and Ca^2+^ (300 nM), which works as a co-agonist to channel opening, as demonstrated in numerous electrophysiological studies^7–11^.

From a structural perspective, AdA is intriguing because this fungal glyconucleotide metabolite mimics InsP_3_ by acting as a full agonist that binds to InsP_3_R1 with ~10-fold greater affinity and ~12-times more potency in opening the channel than InsP_3_^7,8,12^. Previous studies suggest that the 3″,4″-bisphosphate and 2′-hydroxyl groups of AdA mimic the essential 4,5-bisphosphate and 6-hydroxyl of InsP_3_, respectively (Supplementary information, Figure S1a)^6,8,13^. The 2′-phosphate is believed, at least in part, to mimic the 1-phosphate of InsP_3_^6,14,15^. This structural similarity between the two ligands likely accounts for the competitive binding of AdA to the same InsP_3_-binding domains (Supplementary information, Figure S1b,c). However, the molecular basis for the unique properties of AdA is unknown, as is the mechanism of channel opening upon ligand binding.

In this study we collected large data sets of both AdA-InsP_3_R1 and Apo-InsP_3_R1. Due to the potential for partial ligand occupancy, the AdA-InsP_3_R1 map was generated using standard single-particle 3D reconstruction techniques combined with a masked focused classification approach to achieve consistency among the particles used in the reconstruction (Supplementary information, Figures S2, S3; see Materials and Methods). The final maps were of sufficient resolution to enable an unambiguous interpretation of the protein structure (Supplementary information, Figures S2, S4). Map resolvability in both states permitted a backbone trace for ~80% of the entire protein, as well as a full atomistic representation in specific domains (~37.5% of the entire protein) (Supplementary information, Figure S5).

The map resulting from the class of AdA-bound InsP_3_R1 particles exhibited robust density bridging the β-TF2 and ARM1 domains (Figure 1). The location of this density is equivalent to the bound InsP_3_ molecule in the crystal structures of isolated ligand-binding domains (Supplementary information, Figure S6a, c)^16,17^. Therefore, the densities identified at the β-TF2/ARM1 cleft in the AdA-InsP_3_R1 map were assigned to the bound AdA molecule.

**Figure 1.**
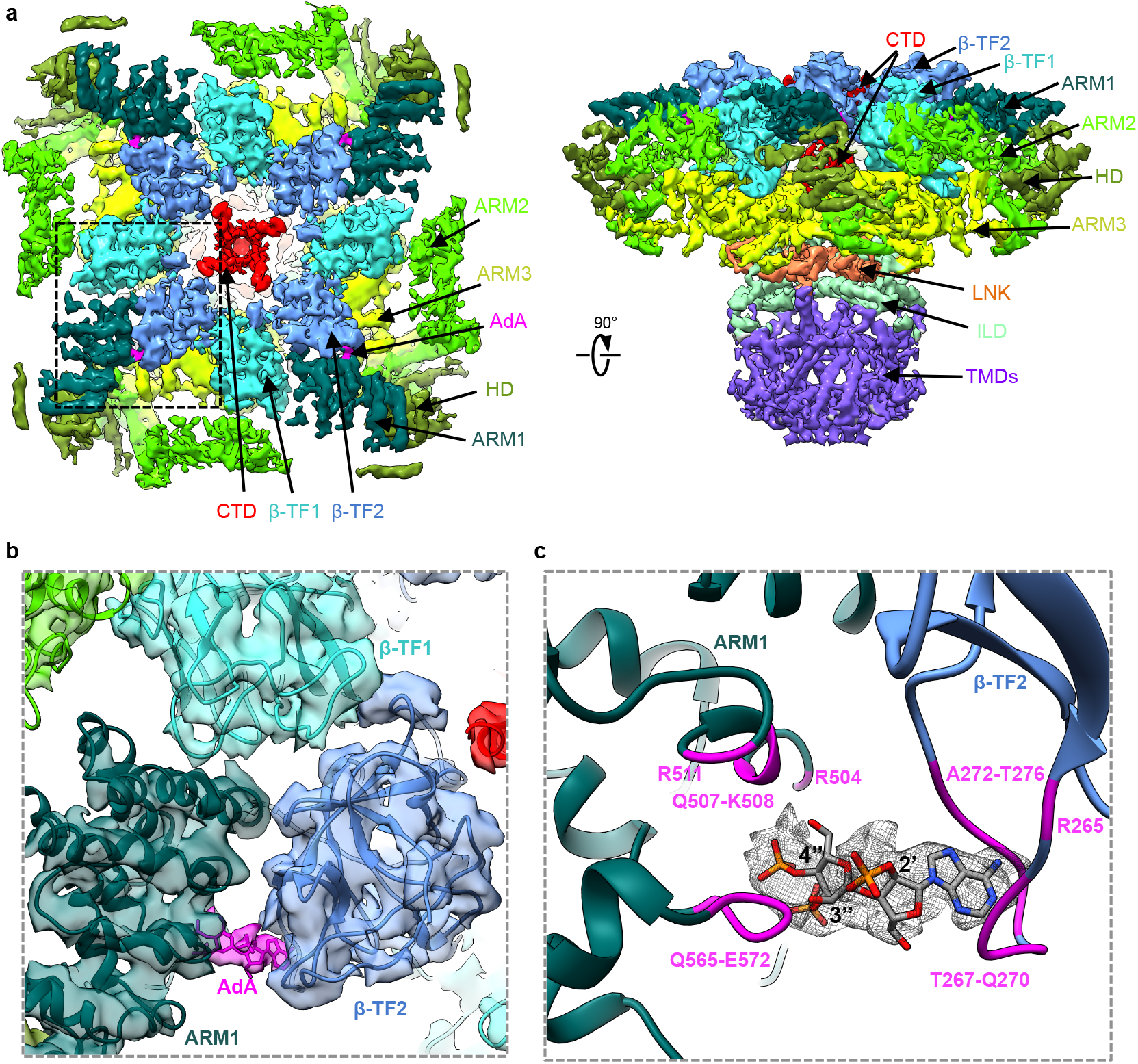
The cryo-EM structure of InsP_3_R1 channel bound with AdA. **a**, The cryo-EM density map of the InsP_3_R1-AdA complex is viewed from cytosol along its four-fold axis (left) and parallel to the membrane plane (right). The map is filtered to 4.1 Å and corrected with a temperature factor of −100 Å^2^. The domains within each protomer are delineated by different colours. Densities corresponding to AdA are coloured magenta. **b**, Zoomed-in view of the AdA density present in the ligand binding pocket. AdA-InsP_3_R1 structure is overlaid on the corresponding density map and coloured by domain; the AdA molecule is fit in the density adjoining the LBDs. **c**, AdA molecule is shown in the InsP_3_R1 binding pocket and is overlaid with densities (mesh) from the difference map (4σ). Residues within 5 Å distance from the docked AdA molecule are labeled and coloured magenta.

The 3D structure of AdA-InsP_3_R1 matches an overall arrangement of the Apo-InsP_3_R1, also determined to 3.9 Å resolution in the present study (Supplementary information, Figures S2, S4, S5): the cytosolic (CY) domains are organized as solenoid scaffolds around a central left-handed four-helix bundle comprised of the C-terminal domains (CTD) from each subunit. The CY and transmembrane (TM) portions are hinged at the cytosolic-membrane interface via the intervening lateral (ILD) and linker (LNK) domains within each protomer of the tetrameric channel assembly (Figure 1a; Supplementary information, Figure S5a, b). Our new structures reveal higher resolution features throughout the entire protein yielding improvements in the domain folds and loop connectivities relative to the previously reported Apo-InsP_3_R1 structure^2^. Many side-chain densities were visible throughout the ten protein domains comprising each protomer (Supplementary information, Figure S5c, d).

### The ligand-binding pocket

The ligand-binding pocket is formed at the cleft between the β-TF2 domain (W226-V435*) and the first two α-helical armadillo repeats of ARM1 (S436-K604). This region can bind InsP_3_ with even higher affinity than the full-length protein when expressed as an independent soluble entity. At the current resolution, the secondary structure elements and overall fold of the LBD are consistent with the known X-ray structures^16,17^. At ~1.5Å RMSD, the final cryo-EM based model is similar to the previously reported X-ray structures, though our model resolves a number of loops that were not previously observed (Figure 1c; Supplementary information, Figure S6c). The AdA molecule was docked in the ligand-binding pocket of the AdA-InsP_3_R1 map, and possible positions were screened based on their visual match to the density (Supplementary information, Figure S6a, b; see Materials and Methods). The final placement of the AdA molecule had the greatest overlap with the bridging density observed between the β-TF2 and ARM1 domains. Based on the docking, AdA appears to be buttressed by two loops, formed by R265-S277 of β-TF2 and D566-Q571 of ARM1, and the ARM1 helix R504-E512 (Figure 1b, c). To adopt a conformation suitable for ligand binding, the β-TF2 and ARM1 loops move toward the ligand, while the R504-E512 helix is relatively unchanged (Figure 2; Supplementary information, Movie S1). While both InsP_3_ and AdA bind to the same site, our study shows that they utilize a different set of parameters to coordinate within the ligand-binding pocket (Figures 1c; Supplementary information, Figure S6c). Specifically, the vicinal 3″- and 4″-phosphate groups of AdA extensively interact with residues of the ARM1 domains, while the adenine ring of AdA interfaces with residues of the β-TF2 domain on the opposite side of the ligand-binding pocket (Supplementary information, Movie S1). These complementary interactions provide the impetus for pulling the two domains towards each other, reducing the interstitial space of the binding pocket by 5 Å in the AdA-InsP_3_R1 structure (Figure 2b). By contrast, the 4- and 5-phosphate of InsP_3_ interact with the β-TF2 and ARM1 domains, respectively, to drive the closure of the InsP_3_-binding cleft (Supplementary information, Figure S6c)^16,17^.

**Figure 2.**
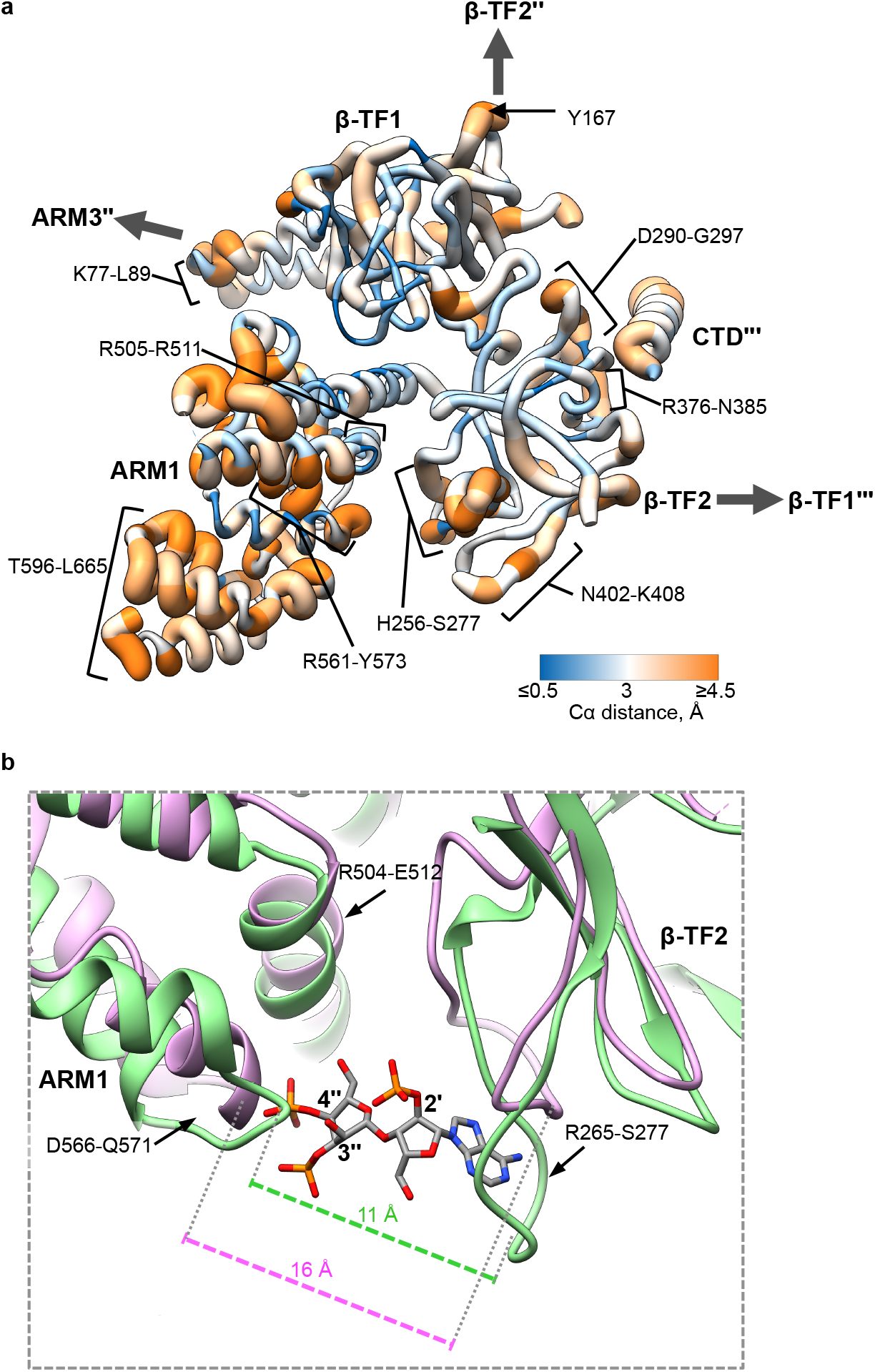
Structural rearrangements in the ligand-binding pocket of InsP_3_R1. **a**, The Cα RMS deviations calculated between Apo- and AdA-bound LBD structures are shown in the AdA-bound LBD structure colour-coded based on RMS deviation - ribbon color/thickness denotes lowest RMS (blue/thinnest) to highest RMS (orange/thickest) deviations. The most variable residues contributing to intra- and inter-domain contacts are labeled. Wide arrows point to the interacting domains of neighboring subunits. **b**, Shown is an overlay of ribbon diagrams of the ARM1 and β-TF2 domains from Apo-InsP_3_R1 (light purple) and AdA-InsP_3_R1 (green). The structure of AdA molecule is colour-coded by element (phosphorous is orange; oxygen is red; nitrogen is blue; carbon is gray) and shown within the binding pocket; 2′-, 3″- and 4″-phosphates are labeled. Distances shown are measured between Cα atoms at the narrowest point within the ligand binding pocket for AdA-bound (between T266 and K569) and Apo-InsP_3_R1 (between G268 and K569).

The electrostatic potential of the ligand-binding pocket is mostly positive, though the β-TF2 ligand binding loop contributes to a small negative electrostatic patch (Supplementary information, Figure 6c). This electrostatic potential of the ligand-binding pocket is complementary to the charge of AdA and may serve as a mechanism to localize AdA: multiple basic amino acids could facilitate ionic interactions with the negatively charged AdA phosphates while the negative electrostatic patch may aid in anchoring the positively charged adenine moiety.

### Rearrangements in the ion conduction pathway

Local resolution estimates and feature visualization indicate that the TM region in both the Apo- and AdA-InsP_3_R1 structures contains better than 3.5 Å resolution information allowing for accurate assignment of side-chains along the ion permeation pathway (Supplementary information, Figures S2f and S5c,d). The domain-swapped arrangement of the six TM helices in InsP_3_R1 unveiled in our previous study^2^ is consistent with a long TM4-5 α-helical linker that would be inserted laterally into the cytosolic leaflet of the membrane (Supplementary information, Figure S7a). The densities on the lumenal side of the TM region are likely attributed to loops connecting TM helices. The extended lumenal TM5-6 loop is structured and harbors the pore helix (P-helix) and the selectivity filter (SF). A previously unassigned density observed between TM1 and TM2 (TM1-2) in both maps, is strikingly similar to the TMx identified in the RyR1 map. While the question of the identity of TMx in RyR1 remains open^18^, TM1-2 density is likely a part of the InsP_3_R protein given its contiguous connectivity with TM2 and that no additional proteins were identified in our InsP_3_R1 preparations^2^.

The ion conduction pore of InsP_3_R1 is shaped by four pairs of the inner TM6 and outer TM5 helices that form a right-handed bundle with a tapering pathway for ion permeation through the membrane (Figure 3a, Supplementary information, Figure S7). In the Apo-InsP_3_R1 structure, the pore is maximally constricted at F2586 and I2590 of the pore-lining TM6 helix. Here, the side-chains form two hydrophobic rings near the cytosolic leaflet of the ER membrane (Figure 3a, b); the minimum distances across the ion conduction pathway at F2586 and I2590 are 4.5 Å and 6 Å, respectively (Figure 3c, d), which will preclude the passage of hydrated cations (*e.g*., Ca^2+^, Na^+^ and K^+^). Thus, our Apo-InsP_3_R1 structure is in a non-conductive closed state with F2586 and I2590 of TM6 serving as the pore gate. This assignment is consistent with our previous cryo-EM studies^2^ and supported by multiple mutagenesis and electrophysiological studies previously demonstrating that mutations in this region abolish channel conductance^19,20^. Hence, conformational rearrangements that lead to pore dilation are necessary to allow for ion conductance of InsP_3_R upon activation.

**Figure 3.**
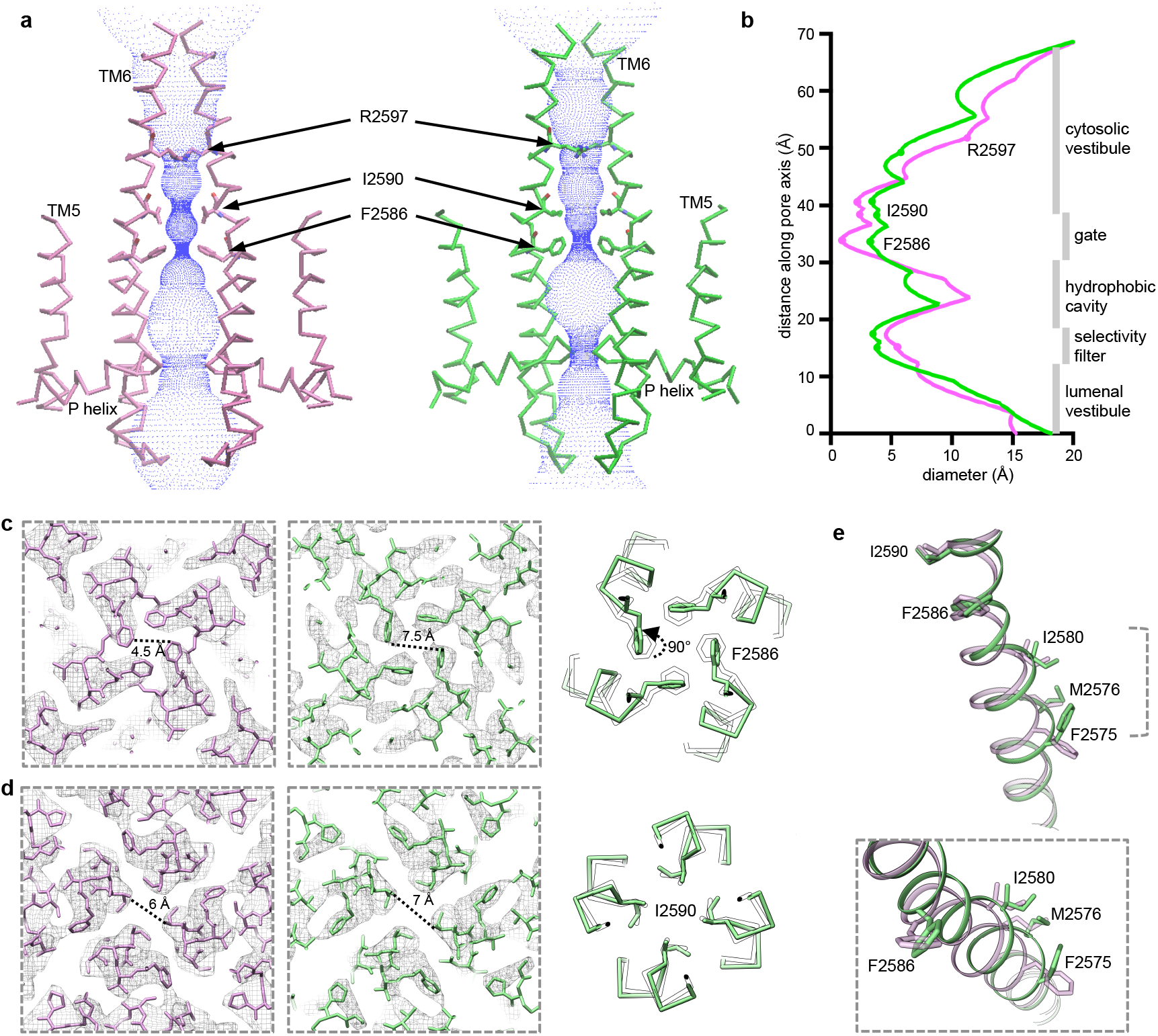
Conformational changes in the ion conduction pathway upon ligand-binding. **a**, Solvent-accessible pathway along the pore mapped using the program HOLE^62^ for Apo- (left) and AdA-bound (right) InsP_3_R1. A series of residues within the ion conduction pathway of the channel pore are labeled. **b**, Comparison of pore diameter for Apo- (light purple) and AdA-bound (green) InsP_3_R1. **c-d**, Sections of density maps perpendicular to the pore at the position of F2586 (**c**) and I2590 (**d**) are shown overlaid with their corresponding models and viewed from the cytosol; Apo-InsP_3_R1 model - light purple, AdA-InsP_3_R1 - green; distances between the sidechains from two opposing TM6 helices are indicated; corresponding side-chain rotations are indicated in the right panels. **e**, Zoomed-in views of the bulge in TM6 seen in AdA-InsP_3_R1 (green) overlaid on TM6 from Apo-InsP_3_R1 (light purple). The lower panel represents a ~40° rotation in view from the upper panel.

Indeed, in the AdA-InsP_3_R1 structure the TM helices exhibit significant structural changes (Supplementary information, Figure S7b). Structural rearrangements of TM helices are characterized by the changes to the central helical axis tilt with respect to their orientations in the Apo-state. The TM6 appears to have three segments (TM6a, TM6b, TM6c) with separate tilt axes that together form the ~70 Å long helix. The change in tilt angle of these three segments is different with the greatest change in tilt axis within the TM6b segment, which contains the residues comprising the gate.

The movements of TM6 result in a bulge away from the 4-fold central axis between I2580 and M2576 and F2575 (Figure 3e). The TM6 conformation may be stabilized by stacking of aromatic residues Y2569, F2573, F2455 observed at the buried interface between TM5 and TM6 helices of the same subunit. This configuration may facilitate a controlled movement of TM6 (Supplementary information, Figure 7b). The TM6 conformational changes observed in the AdA-InsP_3_R1 structure cause the aromatic ring of F2586 to rotate ~90° from the plane normal to the ion conduction pathway. This results in the expansion of the corresponding hydrophobic ring to ~7.5 Å while the hydrophobic seal at I2590 dilates minimally (Figure 3a-d).

Beyond the gate, the ion conduction pathway widens substantially into the cytoplasmic vestibule, where the R2597 residue is positioned (Figure 3a,b; Supplementary information, Figure S8a). Here we propose a network of interactions centered around the R2597 that could play a role in transmitting signals to the gate, in part, through an interaction between neighboring TM6 helices. R2597 is highly conserved among both families of intracellular Ca^2+^ release channels, and mutations R2597A, R2597K and D2591A substantially diminish the channel function, supporting importance of these identified interactions in the cytosolic pore vestibule^20^. Additionally, the TM4-5 linker between the central pore helices and the bundle of peripheral TM1-TM4 helices, is positioned in proximity to the TM6 helix at a TM6-TM6’ interface (Supplementary information, Figures S7, S8a). In the AdA-InsP_3_R1 structure TM4-5 undergoes conformational changes that may alter its interaction with TM6 (Supplementary information, Movie S2). Movement of TM4-5 has been proposed to relieve the steric constriction of the gate allowing it to transition from closed to open conformation ^21,22^.

The central TM5 and TM6 helices provide the structural framework that position the P-helix and the interconnecting loops shaping the lumenal vestibule, which may function as a cation accumulating reservoir positioned before the SF (Figures 3a, 4), a property shared by other cation channels^23^. The SF of InsP_3_R1 is formed by the conserved residues G2546-V2552 (Figure 4a). Notably, the lumenal vestibule in Apo-InsP_3_R1 structure is large enough to accommodate hydrated ions and does not constitute a barrier that precludes ion passage. In the AdA-InsP_3_R1 structure, the P-loop undergoes structural rearrangements resulting in a both a physical constriction and an increase in the electronegative potential in the SF (Figure 4b-d). The separation between Cα atoms from two opposing subunits narrows from 12 Å to 8.5 Å before the entrance to the central hydrophobic cavity. This suggests that in AdA-InsP_3_R1 a Ca^2+^ ion might be able to pass through the filter by, at least partially, displacing its hydration shell, given the diameter of hydrated Ca^2+^ is 8-10 Å^24^.

**Figure 4.**
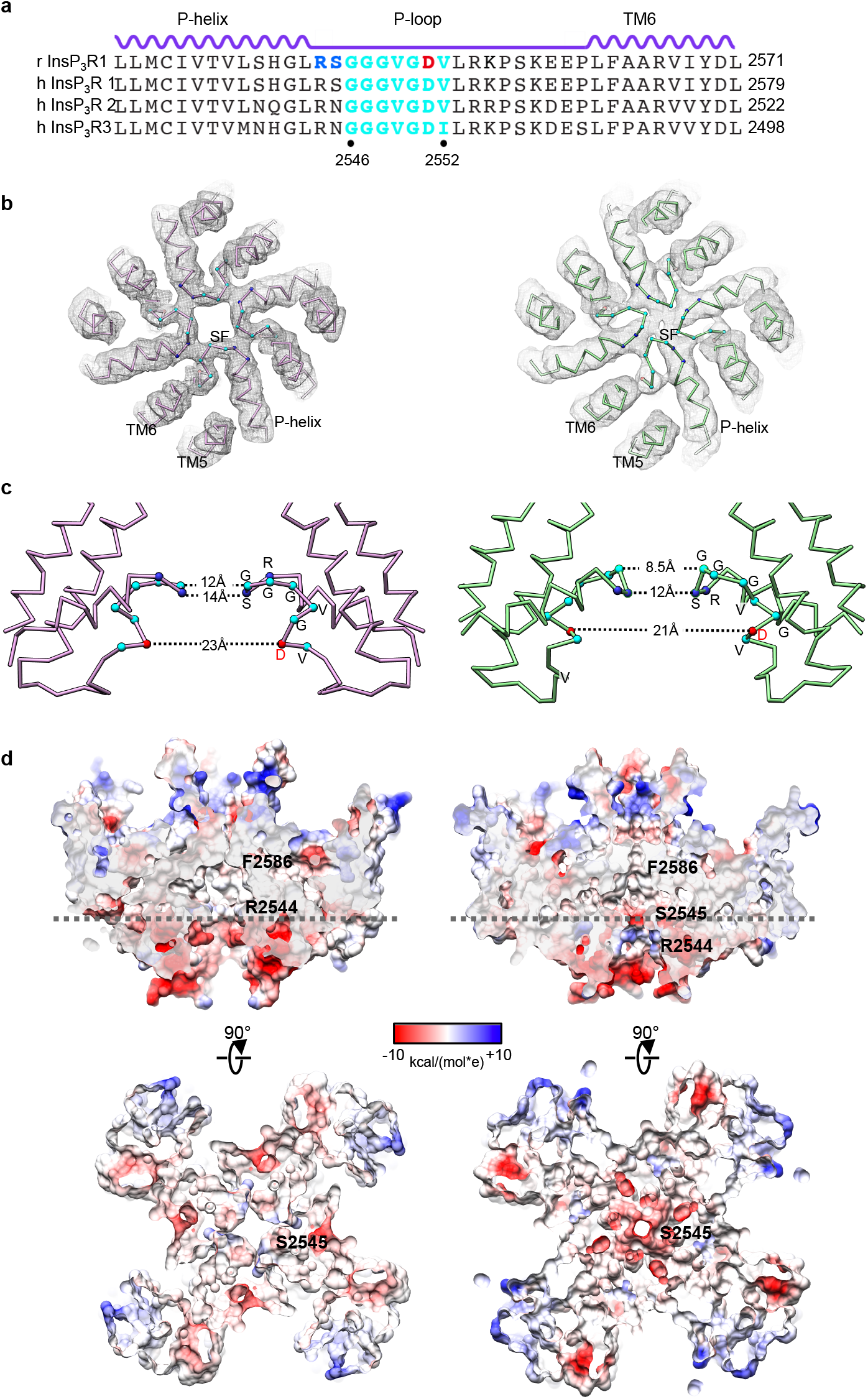
Detailed views of the selectivity filter in InsP_3_R1. **a**, Sequence alignment for the P-loop region of selected InsP_3_R channels; the secondary structure elements are given above sequences; highly conserved residues within the signature sequence of the SF are coloured cyan; residues undergoing significant conformational changes are coloured blue; D2551 within the SF, for which mutations can abolish Ca^2+^ conductance, is colored red; **b**, EM densities with the corresponding models for the SF of Apo- (light purple) and AdA-InsP_3_R1 (green) are viewed from the cytosolic side. **c**, The SF (wire representation) from two opposing subunits is viewed from the side in Apo- (light purple) and AdA-InsP_3_R1 (green); distances between Cα atoms of residues along the SF are indicated; **d**, The surface electrostatic potential in the SF. Top panels show cross-sections along the 4-fold axis through the ion conduction pathway in Apo- (left) and AdA-InsP_3_R1 (right); lumenal entrance at the bottom. Bottom panels - slices through the channel pore perpendicular to the 4-fold axis at the positions indicated with dashed lines in upper panels (viewed from cytosol).

The ligand-induced rearrangements in the SF may confer a weak selectivity of Ca^2+^ over monovalent cations (P_Ca_/P_K_ ~6-8) in InsP_3_R channels visualized in electrophysiological experiments. The modest selectivity of the InsP_3_R1 channel implies that the SF is rather permissive in the presence of other cations, such as Na^+^ or K^+4,25,26^. In the AdA-InsP_3_R1 structure, the positively charged surface preceding the constriction of the SF may serve to focus cations along the ion conduction path via electrostatic repulsion (Figure 4d), and together with the physical and chemical properties of the SF would provide the basis for the screening of permeant cations. The Ca^2+^ ion being more electropositive than Na^+^ or K^+^, would be better coordinated and bind more tightly in the SF. This may prevent entrance of other monovalent cations to the permeation path, thereby providing a plausible mechanism for modest ion selectivity^4,25–27^. Noteworthy, *in vivo* unidirectional Ca^2+^ flux via InsP_3_Rs to the cytosol is largely driven by the high Ca^2+^ concentration in the lumen, ensuring rapid occupancy of the SF. Overall, the importance of rearrangements in the lumenal vestibule is supported by previous mutagenesis within the SF, which showed that D2551A inactivated InsP_3_R1 channel function and D2551E exhibited altered cation selectivity and was nonselective for Ca^2+^ over K^+^, with unchanged ion conductance of both cations^28^.

### Ligand mediated allosteric network

In comparing the AdA- and Apo-InsP_3_R1 structures, striking conformational rearrangements in inter- and intra-subunit interfaces coincident with AdA-binding are evident and likely play a role in propagation of ligand-evoked signals towards the ion conduction pore in order to open the channel gate (Supplementary information, Movies S1, S2). The LBDs (β-TF1, β-TF2 and ARM1) are contained within each InsP_3_R1 subunit and form a triangular architecture comprising an apical portion of the cytoplasmic region, similar to that seen in the earlier crystal structures of the isolated LBDs (Supplementary information, Figure S9)^16,17^. Binding of AdA within the ligand-binding pocket induces global rigid body movements of all three LBDs. Ligand binding in the tetrameric channel results in a 5 Å closure of the cleft between the ARM1 and β-TF2 domains (Figure 2), while β-TF1 acts as a pivot for the concerted movement of ARM1 and β-TF2. The ARM1/β-TF1/β-TF2 angle decreases whereas the β-TF1/β-TF2/ARM1 angle increases (Supplementary information, Figure S9b). These cooperative changes in the relative orientations of LBDs, combined with subtle changes in their internal structure, render the ligand-binding pocket amenable for capturing ligand.

Not restricted to intra-subunit interactions, changes of LBDs within one subunit are propagated to the interfacial elements between adjacent subunits. Notably, the interface between the β-TF1 and β-TF2 domains from the adjacent subunits undergoes conformational changes involving the loop harbring Y167, which is essential for Ca^2+^-release activity (Figures 2a; Supplementary information, Movie S1)^29^. Additionally, the helix–turn–helix motif (S67-L110) of β-TF1 rotates and moves toward the ARM3 domain of the adjacent subunit (Figure 2a, 5a). To compensate, the interfacial loop of ARM3 (S2013-Y2024) undergoes a modest motion combined with a slight structural rearrangement. Of note, one of the ARM3 interfacial loops carries a putative ATP-binding site that has been implicated in the modulation of InsP_3_R gating^30–33^.

**Figure 5.**
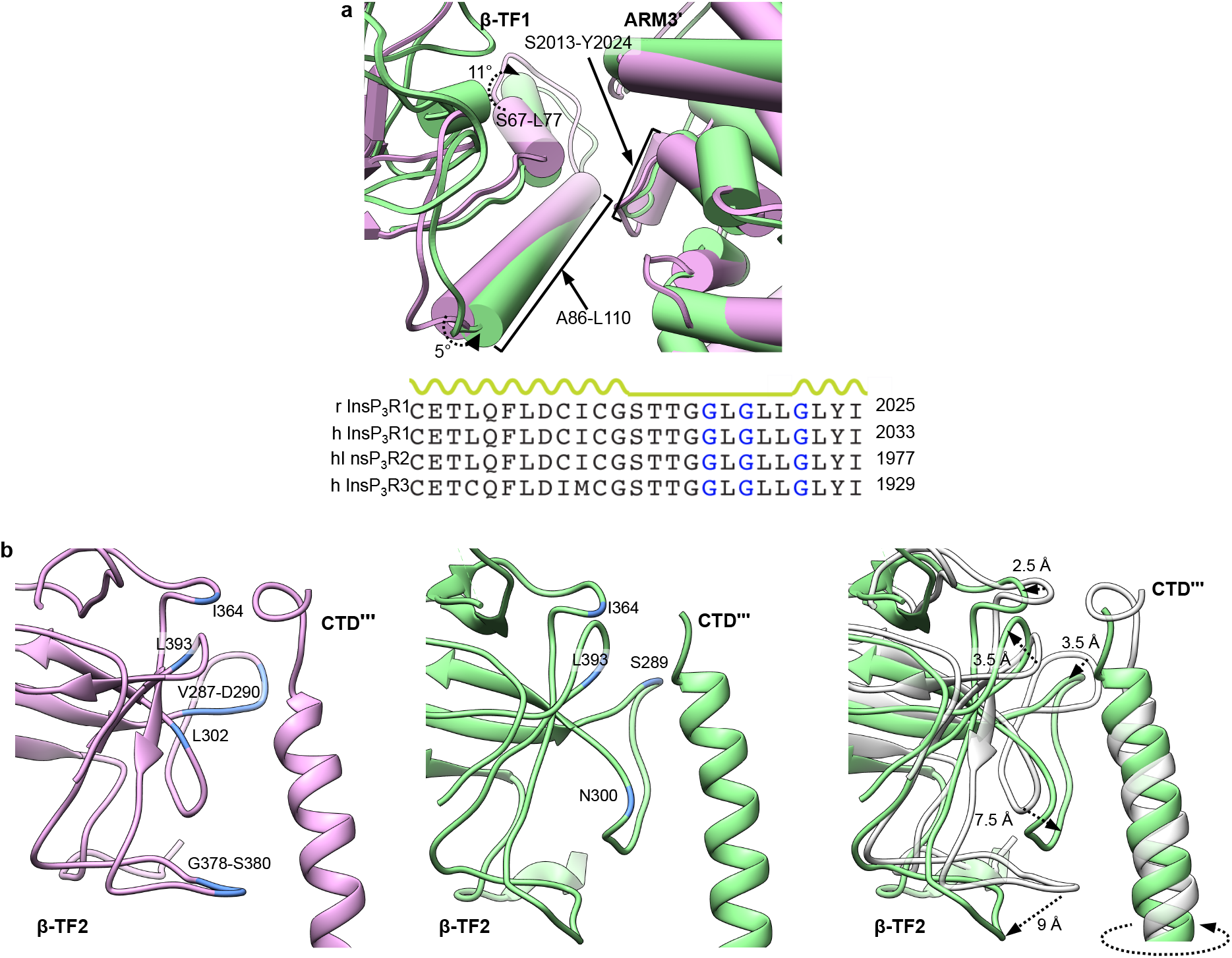
Inter-subunit contacts within cytoplasmic region. **a**, Superimposition of interfaces between β-TF1 and ARM3’ domains in AdA-bound (green) and Apo- (light purple) InsP_3_R1 structures; helices are rendered as cylinders. The lower panel shows the ATP-binding consensus (GXGXXG, indicated in blue) within the ARM3 domain. **b**, Interface between β-TF2 and CTD′″ domains in AdA-bound (green) and Apo- (light purple) InsP_3_R1 structures is shown in view parallel to the membrane plane. Residues within 5 Å of the CTD are coloured blue and labeled. The right panel shows the overlay of AdA- (green) and Apo- (gray) InsP_3_R1 models. Notable structural changes are indicated. In the AdA-InsP_3_R1, the CTD helix exhibits pronounced rotation compared to its position in the Apo-structure.

The β-TF2 domain is located in close proximity to the CTD, located over 2000 amino acids away in the InsP_3_R1 protomer, of the adjacent subunit (Figure 5b; Supplementary information, Movie S1). It follows then that the ligand-evoked structural changes of β-TF2 trigger a pronounced rotation of the CTD helix and this motion propagates through the CTD helix, thereby directly coupling remote but functionally coordinated domains (Supplementary information, Movie S2). Importance of this structural coupling was predicted based in earlier biochemical studies^17,21^. Based on our structures, the ILD/LNK sandwich, which resides near the cytosolic face of the membrane in the tetrameric channel assembly, represents the sole direct structural link between the cytoplasmic and transmembrane domains (Figure 6). The CTD is directly connected to the LNK domain, thus enabling a mechanism for cooperative communication between the ligand-binding and transmembrane domains. Notably, portions of the ILD/LNK sandwich move horizontally away from the four-fold axis following binding of AdA (Supplementary information, Movie S3). Mutations within the first two strands (I2196-E2217) of the ILD domain support the mechanism of signal transduction through the assembly formed by the interleaving ILD and LNK domains at the membrane cytosol interface^34^.

**Figure 6.**
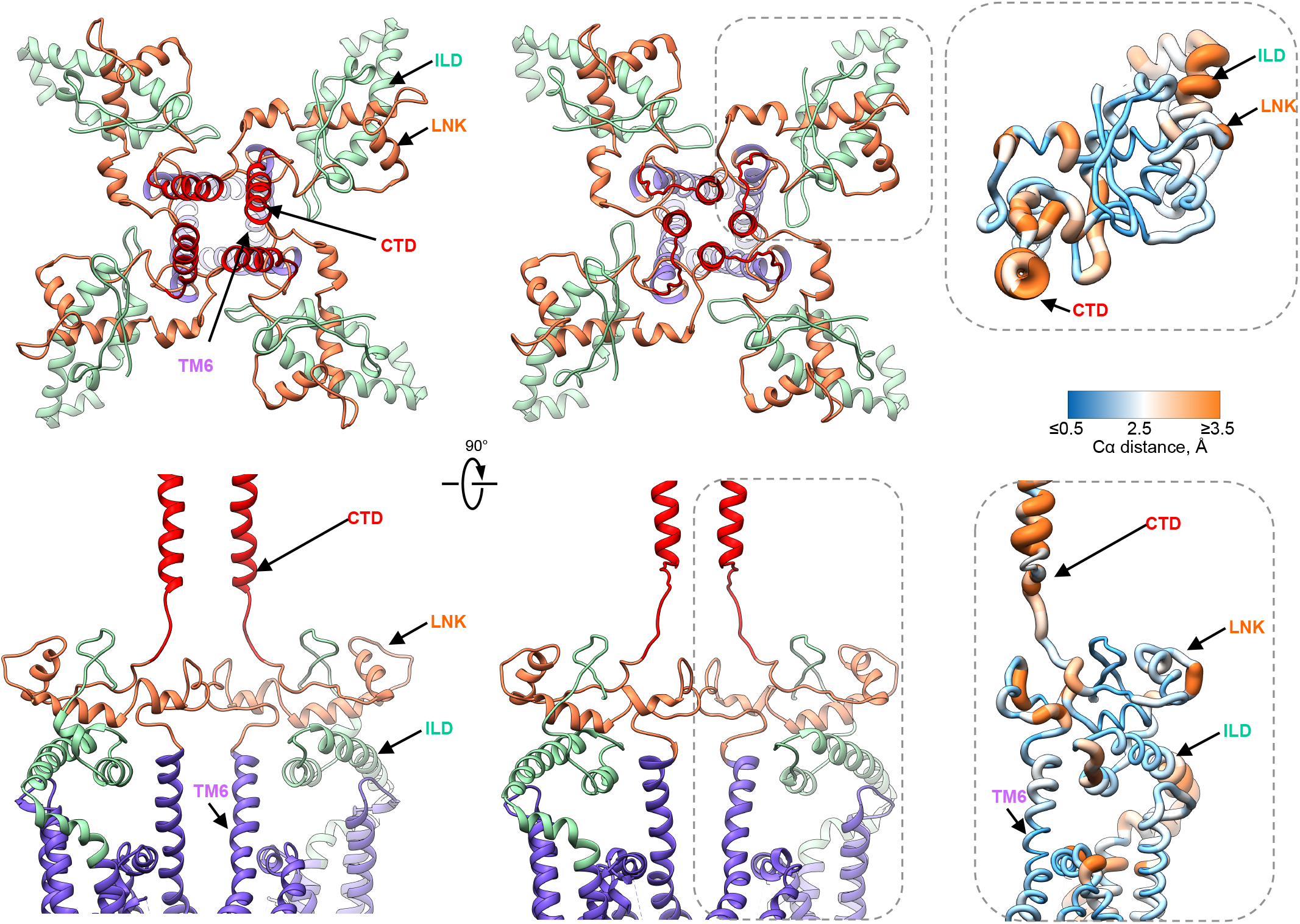
Domain rearrangements at the cytosolic-membrane interface. **a**, Two orthogonal views of domains at the cytosolic-membrane interface in the Apo- (left) and AdA-bound (right) InsP_3_R1 structures (colour-coded by domain). Structural differences are notable in the CTD helices (red), which are connected to the LNK domain (orange). Ligand-triggered conformational changes are ultimately funneled to the TMDs (purple) through the interface comprised of the ILD/LNK sandwich. The lower panels show only two opposing subunits for clarity. Inserts show zoomed-in views (indicated with dashed-lines) of domains from one subunit of AdA-InsP_3_R1 that are colour-coded based on the Cα RMS deviations calculated between Apo- and AdA-structures: ribbon color/thickness denotes lowest RMS (blue/thinnest) to highest RMS (orange/thickest) deviations.

However, the ILD conformation in AdA-InsP_3_R1 is quite different from that observed in the crystal structure of an isolated cytoplasmic portion of the protein, which lacks the molecular constraints provided by the tetrameric assembly of the full-length InsP_3_R1^34^. The putative Ca^2+^-sensor region containing conserved E2101^35,36^ sits at the ARM3-LNK interface that undergoes modest conformational changes in our experimental conditions (Supplementary information, Movie S3). Alignment of 3D structures of two related intracellular Ca^2+^ release channels, InsP_3_R1 and RyR1, shows a conservation of the protein fold within this region, with the latter also containing a structurally conserved putative Ca^2+^-binding site in the same location (Supplementary information, Figure S8b)^3,18^. While the buffer conditions for the AdA-InsP_3_R1 structure include Ca^2+^, the cation concentration was targeted for the channel activation rather than saturation of the Ca^2+^ binding sites, and as a result the densities that could be attributed to Ca^2+^ ions were not observed in the AdA-InsP_3_R1 map.

## Discussion

The present study describes 3D structures of neuronal type 1 InsP_3_R channel with and without the agonist bound. These structures bring into focus molecular features of the ligand binding domains (LBDs) and ion conduction pathway, in which conformational changes are required for activation of the channel gate to enable passage of Ca^2+^ ions.

Conformational sensitivity of InsP_3_R to ligands is a fundamental property of this ion channel underlying its fascinating ability to function as signal transducer. While binding of both AdA or InsP_3_ to InsP_3_R causes activation of the channel, it is conceivable that the differential interactions of ligands within the LBDs would confer the unique properties of the protein that allow of AdA to bind with greater affinity and potency^7,8,12^. This view is supported by earlier studies with synthetic analogues of AdA and InsP_3_^7,15^. Our analysis suggests that the 3″- and 4″-phosphate groups and adenine moiety are important determinants of the increased affinity and high-potency of AdA for InsP_3_R. Our structure reveals that the adenine moiety of AdA interacts with residues of β-TF2 (Figure 1c; Supplementary information, Figure S6c) rather than with the ARM1 as previously proposed^7^. Furthermore, the 2′-phosphate does not appear to interact with the β-TF2 domain, in contrast to the corresponding 1-phosphate of InsP_3_, suggesting that it likely does not contribute to the high-affinity binding of AdA. This is consistent with earlier functional and ligand-binding studies demonstrating that the AdA analogues lacking adenine, but retaining the 2′-phosphate, typically have a binding affinity similar to InsP_3_^7,14^.

Our study supports the concept of long-range conformational changes that propagate from ligand-binding *N*-terminal domains towards the ion-conducting pore through an allosteric network of specific inter- and intra-subunit contacts (Figure 7). This study visualizes binding of AdA at the cleft between the β-TF2 and ARM1 domains that promotes conformational changes in the ligand-binding domains. In turn, this triggers concerted rearrangement of the cytoplasmic solenoid structure organized around the central CTD helical bundle. The interfacial region formed by the ILD and LNK domains undergoes lateral movements that appear to be mechanically connected to pore opening, whereby changes in the ILD/LNK interface generate a force on the TMD via their connecting linkers. As a result, the TMDs undergo rearrangement with the SF adopting an optimal conformation favorable for screening and conduction of cations. The resulting rearrangement in the pore forming helical bundle leads to dilation of the hydrophobic constrictions at the crossing of four TM6 helices.

**Figure 7.**
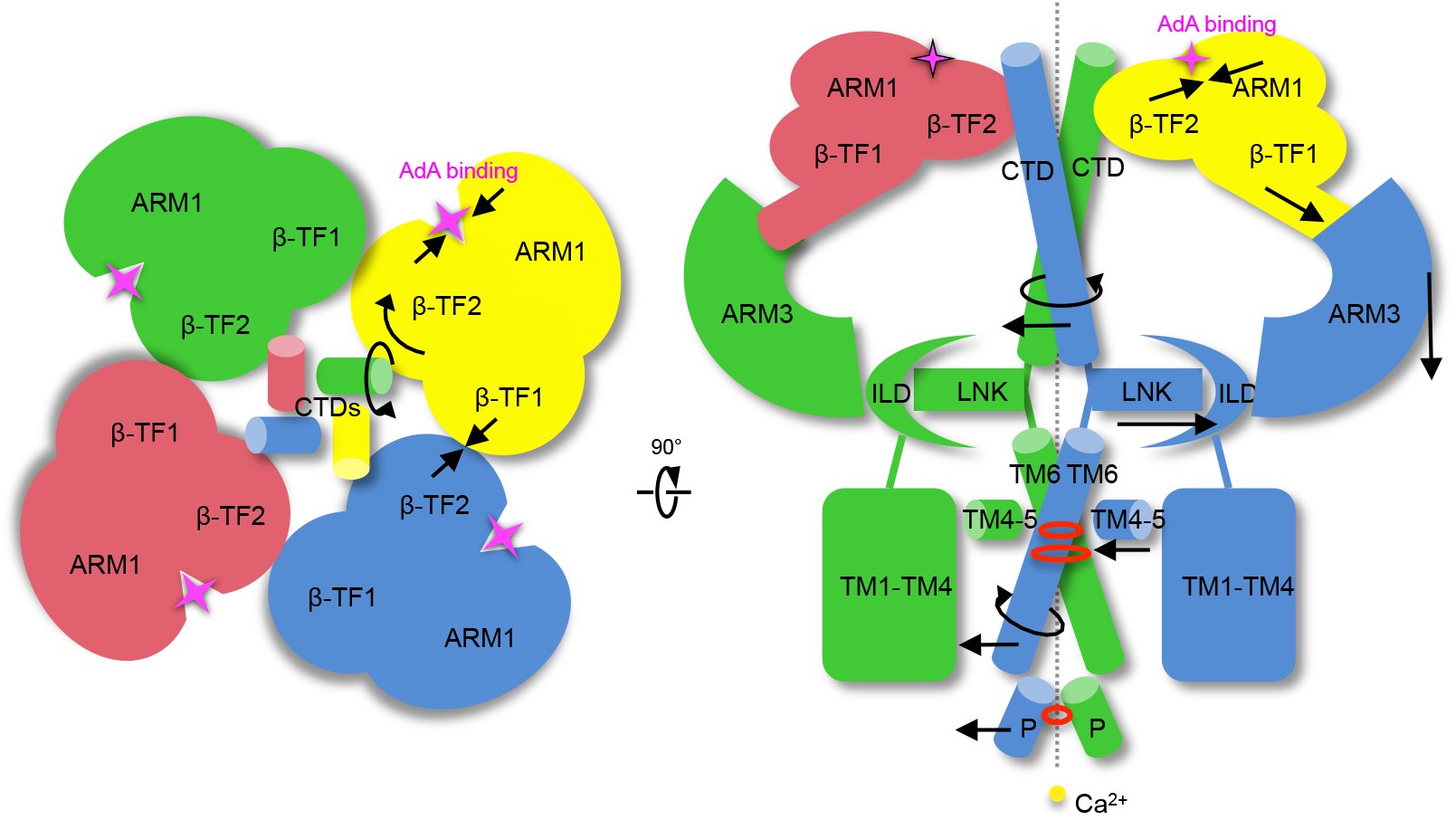
Schematic of ligand-induced conformational changes underlying activation of InsP_3_R1. Top view of the channel along the 4-fold axis from the cytosolic side (left panel). Depicted are the LBDs and CTDs for each InsP_3_R1 protomer coloured by subunit. Section of tetrameric InsP_3_R1 through its 4-fold axis is viewed parallel to the membrane plane (right panel); regions of constriction and SF are indicted with red circles, Ca^2+^ ion - yellow sphere. Ligand-evoked domain motions are indicated with arrows. Presumably, conformational changes evoked by binding of AdA between the β-TF2 and ARM1 domains are propagated via several intersubunit interfaces (β-TF1/β-TF2, CTD/β-TF2 and β-TF1/ARM3) in the cytoplasmic solenoid structure to the ILD/LNK assembly that can exert force directly on the TMDs to open the channel gate.

A surprising finding is that the hydrophobic rings at F2586 and I2590 are not sufficiently dilated in the presented AdA-InsP_3_R1 structure to allow a fully hydrated Ca^2+^ ion to pass through the channel. Nevertheless, the substantial ligand-evoked conformational changes observed in the ion conduction pore suggest that we visualized the channel in an intermediate gating state. There are a few entangled issues related to the complex activation kinetics of InsP_3_R channel that might impact homogeneity of the ligand-bound InsP_3_R1 preparations on an EM grid. Even under the most favorable circumstances the maximum open probability reported for InsP_3_R1 approaches 80% since openings are interrupted by brief closures; the key question is what causes these rapid closures? One set of electrophysiological observations shows that InsP_3_R continuously exposed to InsP_3_ switches to some type of non-conducting state^37^. However, this observation is challenged by other studies^38,39^. While neither satisfactorily accounts for their differences, it may reflect some intrinsic property of InsP_3_R, such as desensitization or inhibition that inexorably follows InsP_3_ binding. Moreover, InsP_3_R1 exhibits ‘modal gating’ where the channel switches between kinetically distinct states even in the presence of saturating concentrations of ligand^40^. This phenomena might reflect the rapid flickering of a gate within a liganded channel rather than ligand-binding events^41^. The recent studies of concatenated InsP_3_R1 raise an additional possibility that the channel opens only when all four ligand-binding sites are occupied^42^. This possibility is somewhat contentious given that it would require substantial increase in InsP_3_ concentration to achieve appreciable activity of InsP_3_R channels *in* vivo^43^. While our structures begin to establish a paradigm for ligand modulated InsP_3_R gating, further high-resolution studies of different ligand-bound states will be necessary to distill these issues.

## Materials and Methods

### Protein purification and ligand-binding assay

All biochemical reagents were obtained from Sigma-Aldrich, Co. unless otherwise stated. Detergent-solubilized InsP_3_R1 was purified from rat cerebellum as described in our earlier studies^2,44^. To assess the ability of purified InsP_3_R1 to bind AdA, we carried out equilibrium-competition ^3^H-InsP_3_ binding assays of cerebellar microsomal membranes and detergent purified receptors in buffer closely representing cryospecimen conditions. For microsomal membranes (100 μg), ^3^H-InsP_3_ (PerkinElmer or American Radiolabeled Chemicals, Inc.) binding (5 nM) was measured in 200 μl of 50 mM Tris-HCl pH7.4, 1 mM EDTA, 100 mM NaCl, 2 mM DTT in the presence of various concentrations of either InsP_3_ or AdA (Calbiochem). After a 5 minute incubation at 4°C, samples were rapidly vacuum filtered through Whatman GF/F filters and washed once with 2.5 ml ice-cold water. Non-specific binding was determined by addition of 10 μM InsP_3_ or 1 μM AdA. For purified InsP_3_R (3-5 μg), ^3^H-InsP_3_ binding (5 nM) was measured in 200 μl of 50 mM Tris-HCl pH7.4, 1 mM EDTA, 100 mM NaCl, 2 mM DTT, 0.4% CHAPS, 0.1 mg/ml BSA in the presence of various concentrations of AdA. After a 5 minute incubation at 4°C, 50 μl of 50 mg/ml γ-globulin and 500 μl of 30% polyethlyenegycol (PEG) 8000 was added. The mixture was incubated at 4°C for 10 minutes and filtered and washed, as described above. Radioactivity was determined by liquid scintillation counting upon addition of 5 ml Ultima Gold scintillation cocktail (PerkinElmer). Non-specific binding was measured in the presence of 1 μM AdA. Graphs and non-linear regression to determine IC_50_ were generated in Prism 7.0 (Graphpad) and each curve is representative of three independent experiments with error bars denoting SEM (Supplementary information, Figure S1b, c).

### Cryo-specimen preparation and cryo-EM imaging

For cryo-EM analysis, the purified InsP_3_R1 (0.1 mg/ml) in 50 mM Tris-HCl buffer (pH 7.4) containing 150 mM NaCl, 1 mM DTT, 0.4% CHAPS, protease inhibitors^44^ was incubated either with 100 nM of AdA and 300 nM of Ca^2+^ (ligand-bound state) or with 1 mM EGTA (Apo-state) for 60 min on ice. Final free Ca^2+^ concentrations were determined using MaxChelator (http://maxchelator.stanford.edu/oprog.htm). Vitrification of the InsP_3_R1 samples was performed as described earlier^2,44^. Images of ice-embedded InsP_3_R1 in the Apo- and ligand-bound states were acquired using Tecnai G2 Polara electron microscope (Thermo Fisher Scientific, Inc.) operated at 300 kV under low-dose conditions. Images were recorded on a K2 Summit direct detector (Gatan, Inc.) in super-resolution counting mode. Data sets were collected at a nominal magnification of 31,000X corresponding to a calibrated physical pixel size of 1.26 Å on the specimen scale corresponding to super-resolution pixel size of 0.63 Å. The dose rate on the camera was 10 electrons/physical pixel/second. The total exposure time of 6 s (for Apo-state) and 7 s (for ligand-bound state) was fractionated into 30 and 35 subframes, respectively, each with 0.2 s exposure time, giving the total accumulated dose of 38 electrons Å^−2^ and 44 electrons Å^−2^ at the specimen plane, respectively. Images were acquired using SerialEM^45^ at a defocus range of 0.8 to 3.5 μm (Supplementary information, Table S1).

### Image processing

Image processing was performed independently using RELION and EMAN2. Both software packages achieved near-atomic resolution in large regions of the structure. However, based on a local resolution assessment performed independently for each map, different domains were better resolved by each software package. To avoid human bias and extract the most information from each reconstruction the final interpreted maps were locally filtered averages of the EMAN2 and RELION maps for both Apo- and ligand-bound states. To combine the two maps, a local resolution filter, based on a windowed FSC local resolution assessment, was performed independently on the two maps. The two locally filtered maps were then averaged together. The local filtration determines the contribution of each map at each resolution in each region of the final composite map, permitting each map to dominate in regions where better self-consistency was obtained during refinement. We can only hypothesize the cause for the differing resolution distribution between the software packages, but it likely relates to the fact that RELION operates purely in Fourier space whereas EMAN uses a hybrid of real and Fourier space operations. Neither method is intrinsically superior, but the two techniques distribute residual noise and artifacts in different ways in the final reconstruction.

### Image processing and 3D reconstruction in RELION

For both Apo-InsP_3_R1 and AdA-InsP_3_R1 datasets, the raw image stacks were binned 2 x 2 by Fourier cropping resulting in a pixel size of 1.26 Å. Each image stack was subjected to motion correction using *‘Motioncorr’* and the motion-corrected frames were summed using frames 2-17 to a single micrograph for further processing (Supplementary information, Figure S2a, b)^46^. We used *‘e2evalimage.py’* in EMAN2^47^ to select 8,450 micrographs from a total of 9,823 micrographs (Apo) and 9,455 micrographs from a total of 14,686 micrographs (ligand-bound) for subsequent processing. 207,914 (Apo) and 191,646 (ligand-bound) particles were selected using *‘e2boxer.py’* (Supplementary information, Table S1). Defocus and astigmatism were determined for each micrograph using CTFFIND3^48^. 144,194 particles of Apo-InsP_3_R1 and 179,760 particles of AdA-InsP_3_R1 were selected after 2D classification and subjected to further 3D classification in RELION^49^. Our previously published map (EMDB #6369) was low pass filtered to 60 Å resolution and used as a starting model for the refinement. Four-fold symmetry was applied during all refinement steps that were performed as described previously^2,3^. In the RELION postprocessing step a soft mask was calculated and applied to the two half-maps before the Fourier shell coefficient (FSC) was calculated. B-factors were estimated (−300Å^2^ and −170Å^2^, for Apo- and AdA-bound maps, respectively) during post-processing procedure in RELION and applied to the map sharpening step. The resolutions for the final 3D reconstructions using standard 3D refinement approach were 3.9 Å for Apo-InsP_3_R1 (from 65,438 particles) and 4.5 Å for ligand-bound InsP_3_R1 (from 179,760 particles) based on the gold standard FSC 0.143 criteria (Supplementary information, Figure S2d)^50,51^. The corresponding particle orientation distributions are shown in Supplementary information, Figure S2c. Local resolution variations were estimated using ResMap^52^ (Supplementary information, Figure S2f). The ligand-bound dataset was further processed using focused classification and 3D refinement strategy as detailed in the following section.

### Focused Mask 3D classification in RELION

To assess structural variability of the AdA-InsP_3_R1 particle population, we performed a 3D classification focused on the LBDs: β-TF1, β-TF2 and ARM1. First we made 4 copies of each particle, one for each of the 4 symmetry-related orientations. Next, a mask with a soft edge extension (*‘relion_mask_create’*) was applied to one LBD region within the initial AdA-InsP_3_R1 density map from RELION (Supplementary information, Figure S3). The modified map with a masked out LBD region from one subunit was re-projected in 2D using the modified euler angles for each particle from the expanded data set. The generated projection images were subtracted from the corresponding particles in the expanded data set to generate a new data set, which includes only the masked region of each particle. The new data set was subjected to 3D classification, without changing orientation, resulting in two populations of the LBD region. The particles from each LBD class were tracked back to the original, unmodified particle images, which yielded 6 subsets of particles^53^. 3D reconstructions were calculated for each of the six subsets and refined without imposing C4 symmetry. We observed density in the putative ligand binding pocket in a pattern implying non-stoichiometric binding. Hypothetically, these subsets correspond to six different occupancies of AdA in the InsP_3_R1 maps: 0-AdA bound, 1 AdA bound, 2 adjacent AdAs bound, 2 opposite AdAs bound, 3 AdAs bound and 4 AdAs bound. However, only the largest class with 3 putative bound AdAs was processed to the highest resolution resulting in the final map of AdA-InsP_3_R1 at 4.1 Å resolution based on the gold standard FSC 0.143 criteria (Supplementary information, Figure S2d,e and Table S1). The remaining classes had poorly resolved features in the ligand-binding pocket and did not exhibit sufficient statistics to proceed with further analysis

### Image processing and 3D reconstruction of Apo-state in EMAN2

The Apo-InsP_3_R1 reconstruction began with 155,060 putative particles. Two initial low resolution refinements were performed to eliminate bad particles using the standard multiresolution quality evaluation in EMAN2 for this purpose, yielding a set of 27,343 particles achieving a better resolution (due to improved self-consistency of the particle set) than the original larger population. This map had an average resolution of 4.5 Å (0.143 criterion following “gold standard” procedures), with significant local variability^54^. This EMAN Apo-map and the Apo-map from RELION were used to produce an initial, incomplete, PDB model. Despite being incomplete, and having mixed quality in different regions, we then took this model, stripped off all sidechains, leaving a simple backbone, then converted this into a density map, which was further subjected to standard phase randomization procedures beyond 7 Å resolution. With these two measures: removing sidechains and phase randomizing, we are confident that any high resolution details emerging in the final model must be produced by the data, not model bias. This initial model was refined using all particles from images less than 2.4 μm underfocus (139,183 particles), which after bad particle elimination left a set of 100,615 particles used in the final refinement. The final map from this process had an overall resolution of 4.3 Å, ranging from 3.5 Å resolution in the TM domain to ~6 Å resolution in the periphery. It is interesting to note that despite going from 27,343 included particles to over 100,000 included particles, the measured resolution changed only very slightly. This demonstrates that our procedures for identifying the particle subsets with the strongest high resolution contribution is reliable (Supplementary information, Figure S2d and Table S1). In many cases eliminated particles do not disagree with the structure they are being excluded from, but simply reiterate the structure at resolutions where the structure is already well resolved, and may not contribute significantly at higher resolution, due to experimental image quality issues. In such cases eliminating particles will have no real impact on the structure at all. Of greater concern is the possibility that the eliminated particles exhibit an alternative, but equally valid, conformation of the assembly. The key is that we do not claim that the high resolution structures represent the single conformation of the assembly in solution, but simply that this represents one dominate self-consistent population. With the level of compositional and conformational variability we believe exists in this system either dramatically improved high resolution contrast or at least an order of magnitude more particles would be required and future studies will aim to obtain such parameters.

### Image processing and 3D reconstruction of AdA-InsP_3_R1 in EMAN2

The full set of 179,730 putative particles was refined using one of the Apo-state maps as an initial model. Many of these particles were known to be false positives or have significant image quality issues. Following standard refinement protocols in EMAN2, the Apo-InsP_3_R1 starting map was independently phase randomized past 10 Å resolution for the even/odd refinements. A standard refinement targeting 5 Å was performed, and the 76,015 particles most self-consistent with this reconstruction was retained for a second refinement targeting 3 Å resolution. Again the most self consistent AdA-InsP_3_R1 particles were retained (33,680) and a third refinement was done. Since we expect only fractional AdA binding, competitive refinement focused on the TM domain was performed between this AdA-InsP_3_R1 map and the Apo-InsP_3_R1 map. This refinement used the 76,015 particle set and produced the population of particles most consistent with AdA-InsP_3_R1 binding and least consistent with the Apo-state (38,405 particles). Since partial AdA occupancy is also possible within each InsP_3_R1 tetramer, this population will still not be perfectly homogeneous, but further classification did not improve resolution due to the small number of particles remaining in each population. These particles were then refined to produce the final EMAN2 AdA-InsP_3_R map, which had an overall resolution of 4.2 Å ranging from 3.6 Å in the TM domain to 8 Å in some peripheral domains.

## Model building

A new Apo-InsP_3_R1 model was constructed directly from the RELION density map using a modified version of our *de novo* modeling protocol. The X-ray structures (PDB IDs: 3UJ4, 3T8S) for the LBD of InsP_3_R1 were first fit to the density. Starting at the end of the LDB structure and using our previous Apo-InsP_3_R1 structure (PDB ID: 3JAV) as a roadmap, we rebuilt the Apo-model to be consistent with the higher resolution density features observed. Refinement of the Apo model was performed using Phenix on the composite Apo map (default parameters of *‘phenix.real_space_refine’*)^55,56^.

To construct the AdA-InsP_3_R1 model, the Apo model was first divided into the 10 separate domains and fit to the RELION AdA-InsP_3_R1 density map. Each domain was then flexibly fit using a combination of FlexEM^57^ and Phenix real space refinement with default parameters. Fit-to-density scores and differences between the original model and the flexibly fit model were computed in UCSF Chimera to identify regions (CC < 0.5 and/or > 3Å RMSD) needing potential further refinement. Once identified, the map and flexibly fit models were imported into Coot and manually refined to optimize both fit to density and model stereochemistry^58^. The newly refined domain models served as anchor points for extending the models. Once the domain models were complete, the individual domains were re-combined into a single model. Iterative automated real space refinement and manual model refinement was then performed with Phenix and Coot.

Initial rounds of Phenix refinement enforced “good” secondary structure elements, rotamers and Ramachandran angles; subsequent refinements saw a gradual relaxation of these parameters. A final series of refinements were performed with the complete AdA-InsP_3_R1 tetramer using the composite AdA-InsP_3_R1 map. Model statistics, including map-model FSCs, fit-to-density, rotamer outliers and ramachandran outliers were monitored during each iteration (Supplementary information, Figure S4b; Table S1). The model was considered final once these statistics converged. In both the Apo- and AdA-bound models, full atomistic models were maintained for regions where the majority of sidechain densities were present, including the LBD, ILD, LNK, TM and CTD domains. While some sidechain densities were observed and modeled in the remaining domains, the final model in these domains were rendered only as mainchain atoms. The ligand densities were visualized by subtracting the structure-factor normalized Apo-map from the final AdA-bound map using ‘vop subtract’ and contoured to 4 σ in UCSF Chimera (Figure 1c). The location of the densities are consistent with the known InsP_3_ binding site. Molecular docking computations of AdA with the ligand-bound InsP_3_R1 model were performed using Autodock Vina^59^ followed by evaluation of the docking positions match to the density observed between the β-TF2 and ARM1 domains (Supplementary information, Figure S6a, b). The final position of AdA gave the highest correlation fit to densities and was a top-scoring binding mode.

Once the models were completed, local map-model FSCs were calculated using EMAN2 (Supplementary information, Figure S4b). Reported model statistics were generated with Phenix and Molprobity^60^ (Supplementary information, Table S1). Map-model visualization and analysis was performed in Coot and UCSF Chimera. Interfaces described were identified with PDBSum^61^ and HOLE^62^.

## Accession codes

Cryo-EM density maps of InsP_3_R1 have been deposited in the Electron Microscopy Data Bank (http://www.ebi.ac.uk/pdbe/emdb/) under accession codes EMD-XXXX, EMD-XXXX, EMD-XXXX, EMDB-XXXX, EMDB-XXXX and EMD-XXXX. Corresponding atomic coordinates have been deposited in the Protein Data Bank (http://www.rcsb.org/pdb) under accession codes XXXX and XXXX.

## Footnote

*The residue numbering is given according to the primary sequence with the GenInfo Identifier (GI) code 17380349, which includes the first methionine.

## Acknowledgements

We are very grateful to David Yule for impactful discussions of presented results and suggestions on the manuscript. We thank Wah Chiu for providing access to NCMI resources at Baylor College of Medicine. This work was supported by grants from the National Institutes of Health (R01GM072804, R21AR063255, R21NS106968, R01GM080139, P41GM103832, American Heart Association (16GRNT2972000), Muscular Dystrophy Association (295138) and National Science Foundation (DBI-1356306). We gratefully acknowledge the assistance and computing resources provided by the Center for Computational and Integrative Biomedical Research of Baylor College of Medicine and the Texas Advanced Computing Center at the University of Texas at Austin in the completion of this work.

## Authors Contributions

IIS conceived the project; GF, ABS, MRB and IIS purified and characterized the protein; GF, ZW and IIS performed cryo-EM experiments, including cryospecimen preparation and data acquisition; GF, ABS, ZW and SJL analysed data; MLB and MRB built the atomic models; MRB, MLB and IIS interpreted the models; GF, MLB and MRB prepared supporting movies; IIS and MRB have written a manuscript with contributions from all authors.

## SUPPLEMENTARY INFORMATION

**Figure S1. Comparison of AdA and InsP_3_R1. a**, Chemical structure of AdA and InsP_3_ coloured-coded by element (phosphorous is orange; oxygen is red; nitrogen is blue; carbon is gray). Competitive ^3^H-InsP_3_ radioligand binding for rat cerebellar microsomal membranes (**b**) with IC50 for AdA = 7.9 ± 2 nM and InsP_3_= 127 ± 10 nM and detergent solubilized, purified InsP_3_R1 protein (**c**) with an IC50 for AdA = 7.5 ± 2 nM. Shown is one representative curve for each from n=3 experiments.

**Figure S2. Single-particle Cryo-EM analysis of Apo- and AdA-InsP_3_R1. a**, Representative 300 keV electron image of InsP_3_R1 particles vitrified in the presence of activating ligands, AdA and Ca^2+^. **b**, Fourier transform of image shown in (a). **c**, Euler angle distribution of particle orientations in the final refinement rounds in RELION. Each view is represented by a sphere, for which the size is proportional to the number of particles in a given orientation. Top panel: Apo-InsP_3_R1 reconstruction; middle panel: consensus AdA-InsP_3_R1 reconstruction; bottom panel: AdA-InsP_3_R1 reconstruction after 3D focused 3D classification (Material and Methods, Supplementary information, Figure S3). **d**, FSC curves for the cryo-EM 3D reconstructions. The resolution was estimated using the gold-standard FSC 0.143 criterion^50,51^. **e**, The gold-standard FSC curves for the cryo-EM maps generated without imposing c4 symmetry from the classes resulted from 3D focused classification (Materials and Methods; Supplementary information, Figure S3). **f**, The cryo-EM density maps of Apo-InsP_3_R1 (upper panels) and AdA-InsP_3_R1 (lower panels) are colour-coded based on ResMap (see Materials and Methods). The maps are viewed parallel to the membrane plane (left panels); the density slabs coincident with the 4-fold axis (indicated with dashed boxes in left panels) are shown in right panels.

**Figure S3. Workflow for 3D reconstruction of AdA-InsP_3_R1**. Signal-subtracted 3D classification was performed to resolve heterogeneity in the ligand-binding pocket (see Materials and Methods).

**Figure S4. Composite cryo-EM density maps of InsP_3_R1. a**, Isosurface rendering of the composite cryo-EM density maps for Apo- (left panel) and AdA-InsP_3_R1 (right panel) (see Materials and Methods); the maps are viewed parallel to the membrane plane with the cytosolic regions facing up. **b**, The FSC plots obtained for the final models (solid lines) and for the TM domains (dashed lines) when compared to the corresponding composite cryo-EM maps.

**Figure S5. Representative cryo-EM densities. a**, The cryo-EM density map for AdA-InsP_3_R1 is overlaid with the model; shown are two opposing subunits. The domains are colour-coded and labeled according to Fan, *et al*.^2^. **b**, Models for one subunit of Apo- (light purple) and AdA-InsP_3_R1 (green) are overlaid. **c-d**, Representative cryo-EM densities for selected regions are overlaid with corresponding models for AdA-(c) and Apo-InsP_3_R1 (**d**).

**Figure S6. Identification and characterization of the AdA binding pocket. a**, Cryo-EM density maps for the ligand-binding pocket in the Apo- (light purple) and AdA- (gray mesh) are overlaid; Apo-InsP_3_R1 is depicted with ribbon model coloured by domain. The bridging density visualized between the ARM1 and β-TF2 domains in AdA-InsP_3_R1 map is marked with red arrowhead. **b**, Several candidate positions for docking the AdA molecule generated using AutoDock Vina^59^(left panel) are displayed within the AdA-InsP_3_R1 LBD structure (green). The right panel shows the final AdA molecular docking and its fit to the difference map density. **c**, Structures of isolated InsP_3_-bound LBDs compared with the AdA-bound InsP_3_R1 cryo-EM structure. Top panels: AdA-InsP_3_R1 (green ribbon), middle panels: 3UJ0 (tan ribbon); panels: 3T8S (gray ribbon). InsP_3_ and AdA are colour-coded by element: (phosphorous - orange; oxygen - red; nitrogen - blue; carbon - gray; phosphates are labeled as indicated in Figure S1a). The right panels show the surface electrostatic charges calculated for the corresponding ligand binding pockets.

**Figure S7. Cryo-EM density map for the TM region of Apo-InsP_3_R1. a**, Two views of the Apo-InsP_3_R1 cryo-EM density map for the TM1-TM6 helices and interconnecting loops of two opposing subunits; viewed parallel to the membrane plane with the lumenal side facing down. **b**, Superimposition of TM helices from one subunit of Apo (light purple) and AdA-bound (green) structures; the helices are depicted as cylinders in two views: parallel to the membrane plane (left, cytosolic side up), and rotated 70° (right, viewed from the cytosol). Changes in a rigid body tilt for each TM helix in AdA-InsP_3_R1 with respect to the orientation in the Apo-state are indicated.

**Figure S8. Inter- and intra-subunit contacts within the pore region. a**, Residues on the neighboring TM6 helix within 5 Å of R2597 are shown with sidechains. The lateral membrane-associated TM4-5 is located in close proximity to the TM6-TM6’ interaction site, whereby each TM4-5 helix is positioned to interact with the TM6 helix from the same subunit. Conformational changes observed within this region upon ligand binding may serve to communicate with the gate. **b**, Zoomed-in view of the putative Ca^2+^ sensor region in ARM3 domain of AdA-InsP_3_R1 (colour-coded by domains) that is superimposed with the same domain in the Apo-InsP_3_R1 (light purple; left panel). Residues that may play a role in Ca^2+^ binding based on a structure-based sequence alignment with RyR1^3^ are labeled.

**Figure S9. Comparative analysis of AdA- and InsP_3_-bound ligand binding domains. a**, The triangular arrangement of ligand-binding domains within tetrameric InsP_3_R structure: AdA-InsP_3_R LBD (green), 3UJ0 (gray) and 3T8S (tan). **b**, Angular relationships and relative distances between β-TF1, ARM1 and β-TF2 domains in Apo- and AdA-bound structures of InsP_3_R1 (upper panels) and in the Apo- and InsP_3_-bound crystal structure of isolated LBD (3T8S; lower panels) as estimated based on the center of mass for each domain. It is noticeable, that AdA ligand binding results in a greater domain closure between ARM1 and β-TF2 than InsP_3_. This results in a decrease in the angular relationship of ARM1/β-TF1/β-TF2 domains while the β-TF1/β-TF2/ARM1 angle increases.

**Supplementary Table S1. Summary of Cryo-EM data collection, image processing, 3D reconstruction and model statistics**.

### Supplementary Movies

**Movie S1. Structural rearrangements of cytosolic domains in InsP_3_R1 upon ligand-binding**. Animated Movie shows a morph between Apo- and AdA-InsP_3_R1 models; the LBDs and CTDs are first coloured by subunit, then colour-coded by domain, followed by a zoomed-in view of one LBD and CTD.

**Movie S2. Ligand-evoked global conformational changes in tetrameric InsP_3_R1**. This animated Movie shows a morph between Apo- and AdA-bound models for one InsP_3_R1 protomer colour-coded by domain, while the other subunits are statically depicted in the Apostate and coloured white. The channel is shown in a side view parallel to the membrane plane.

**Movie S3. Structural rearrangements at the interface between the CY and TM regions**. This animated Movie shows the morph between tetrameric Apo- and AdA-InsP_3_R1 models at the CY/TM interface; the model is colour-coded by domain and viewed along the four-fold axis from the cytosol.

